# The genetic architectures of vine and skin maturity in tetraploid potato

**DOI:** 10.1101/2022.04.04.486897

**Authors:** Maria V. Caraza-Harter, Jeffrey B. Endelman

**Affiliations:** Department of Horticulture, University of Wisconsin-Madison, Madison, WI 53706, USA

## Abstract

Potato vine and skin maturity, which refer to foliar senescence and adherence of the tuber periderm, respectively, are both important to production and therefore breeding. Our objective was to investigate the genetic architectures of these traits in a genome-wide association panel of 586 genotypes, and through joint linkage mapping in a half-diallel subset (N = 397). Skin maturity was measured by image analysis after mechanized harvest 120 days after planting. To correct for the influence of vine maturity on skin maturity under these conditions, the former was used as a covariate in the analysis. The genomic heritability based on a 10K SNP array was 0.33 for skin maturity vs. 0.46 for vine maturity. Only minor QTL were detected for skin maturity, the largest being on chromosome 9 and explaining 8% of the variation. As in many previous studies, *S. tuberosum Cycling DOF Factor 1 (CDF1)* had a large influence on vine maturity, explaining 33% of the variation in the panel as a bi-allelic SNP and 44% in the half-diallel as a multi-allelic QTL. From the estimated effects of the parental haplotypes in the half-diallel and prior knowledge of the allelic series for *CDF1*, the *CDF1* allele for each haplotype was predicted and ultimately confirmed through whole-genome sequencing. The ability to connect statistical alleles from QTL models with biological alleles based on DNA sequencing represents a new milestone in genomics-assisted breeding for tetraploid species.

## INTRODUCTION

Maturity is very important for plant breeding and production and can refer to several, related concepts (Bruns 2007). Perhaps the most common meaning is the time interval between planting and harvest, such that varieties with early maturity can be harvested before those with late maturity. The traits that determine when a crop is ready for harvest vary across species. Although potato (*S. tuberosum*) tubers are not true seeds, they still develop toward a condition of maximum physiological dormancy (Struik and Wiersema 1999). Several maturity traits are used to describe the changes that occur during this process (Bussan et al. 2009). Although these traits can refer to a time interval, more commonly they describe a range of phenotypic variation at one time point. Vine maturity refers to senescence of the foliage and has been traditionally measured based on a visual rating; more recently, remote sensing has been used (Sankaran et al. 2015; Colwell et al. 2021). Chemical maturity refers to the concentration of tuber sucrose, which decreases in late development as starch and dry matter levels increase (Sowokinos 1978).

Skin maturity, which is commonly called skin set, refers to adherence of tuber skin to the underlying tissue. Anatomically, skin is the phellem (outer) layer of the potato periderm, which also contains the phellogen (middle) and phelloderm (inner) layers (Reeve et al. 1969).

Phellogen is meristematic tissue, adding cells of suberized phellem to the outside and phelloderm to the inside during tuber growth. Skinning (Figure 1) occurs when the phellem is subjected to shear stress and separates from the phelloderm due to fracture of the phellogen (Lulai and Freeman 2001). Skinning reduces the marketability of tubers and their storability, by compromising the barrier to water loss and pathogen entry. Skin maturity can be quantified based on the percent of the tuber surface without skin, after exposure to the stresses of harvest and grading (Caraza-Harter and Endelman 2020). Another approach is to measure the torque at which skinning occurs, using a handheld torsion meter (Halderson and Henning 1993; Lulai and Orr 1993).

**Figure 1.**
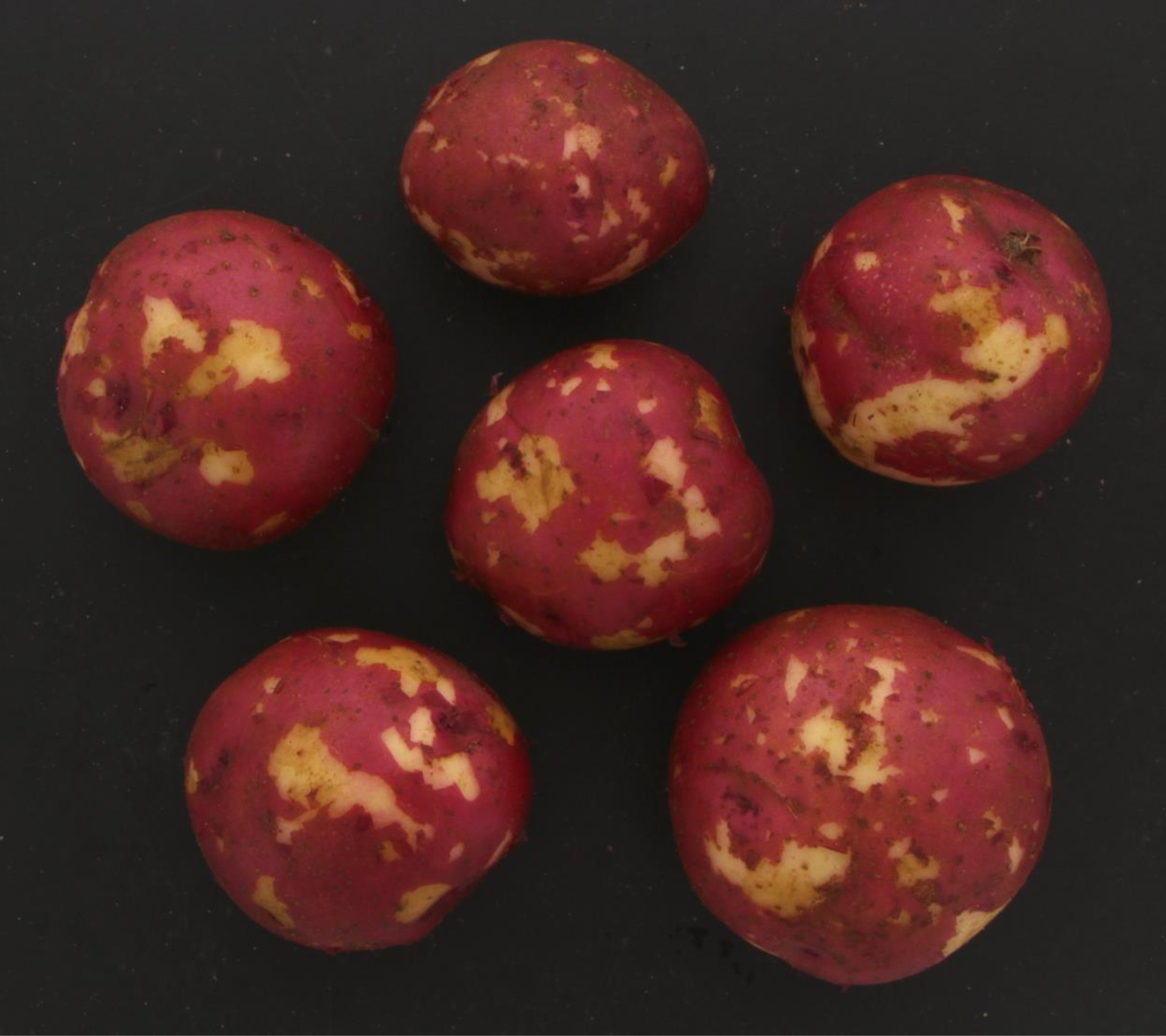
Skinning injury on a red-skin potato tuber. The clone pictured is W10209-7R, one of the checks in the 2017 field trial. Based on image analysis, 18% of the tuber surface was skinned.

Little is known about the genetics of skin maturity, in part because it is rarely measured in breeding populations. Some progress has been made by studying patterns of gene expression in select varieties (Neubauer et al. 2013; Vulavala et al. 2019), which indicates it is a complex trait. Our understanding of the relationship between skin and vine maturity is primarily based on research geared toward crop management, not breeding. Management actions that hasten foliar senescence, such as chemical desiccation or reduced fertilization, generally lead to better skin maturity at a fixed days after planting (Murphy 1968; Bethke and Busse 2010). As such, the predominant conceptual model is that skin maturation follows vine maturation. In contrast to skin maturity, vine maturity is routinely measured in breeding and has been the subject of many QTL studies (e.g., Collins et al. 1999; Bradshaw et al. 2004; Uitdewilligen et al. 2013).

Most studies of vine maturity identify a major QTL on potato chromosome 5, which Kloosterman et al. (2013) demonstrated was a potato homolog of the *A. thaliana* gene *Cycling DOF Factor 1* (*CDF1*). *S. tuberosum* CDF1 promotes expression of florigen (SP3D) and tuberigen (SP6A), which are key signals for the initiation of flowering and tuberization, respectively (Navarro et al. 2011; Abelenda et al. 2014). The connection between this pathway and foliar senescence (i.e., vine maturity) is not well understood. In addition to the DOF (DNA-binding One zinc Finger) domain, the ancestral (i.e., wild-type) *S. tuberosum* CDF1 contains C-terminal domains that interact with the proteins GI and FKF1 in a daylength-dependent manner (Kloosterman et al. 2013; Gutaker et al. 2019). Under short days, such as those found in the Andean region where potato was first domesticated, studies of *A. thaliana* show FKF1 expression lags behind GI (Sawa et al. 2007). Under long days, their expression becomes more synchronized, leading to diminished activity of wild-type CDF1.

Three alleles of potato *CDF1* that code for truncated proteins have been discovered (Kloosterman et al. 2013; Gutaker et al. 2019). These alleles are numbered 2–4 (wild-type alleles are collectively designated type 1), and all contain an insertion in the same location, creating a premature stop codon. *CDF1.3* has an 865 bp insertion from a putative transposon, while *CDF1.2* and *CDF1.4* have different 7 bp insertions. These truncated CDF1 variants are missing the FKF1 binding domain, which leads to enhanced CDF1 activity and early vine maturity under long days. The large size of the insertion in *CDF1.3* also disrupts a long, non-coding RNA named *FLORE*, which is anti-sense to the *CDF1* transcript (Ramírez Gonzales et al. 2021).

The present study was initiated to study the quantitative genetics of skin maturity and its relationship to vine maturity in breeding populations. Since beginning the study, new software has been developed to characterize the function of haplotypes in tetraploid, multiparental populations (Zheng et al. 2021; Amadeu et al. 2021). Sequencing technology has also advanced, reducing the cost and time needed to determine the sequence and dosage of haplotypes. Using these new tools, the original scope of the study was extended to investigate the sequence-function relationship between *CDF1* and maturity.

## MATERIALS AND METHODS

### Germplasm, genotyping, genomics, and phenotyping

The study population consisted of two consecutive cohorts from the red-skin potato breeding program at the University of Wisconsin-Madison. The 2017 cohort consisted of 127 clones, of which 47 were selected (based on overall merit) for a second year of evaluation with the 2018 cohort of 459 clones, for a total population size of 586. The 2018 cohort was much larger than normal because minimal selection was practiced in the first two field generations, to prepare for the present study. Breeding lines were genotyped with version 3 of the potato SNP array (Felcher et al. 2012; Vos et al. 2015), and genotype calls were made using R package fitPoly (Voorrips et al. 2011; Zych et al. 2019). Each marker data point is the estimated dosage (0–4) of the B allele for the sample. There were 10,185 SNPs after removing markers for which fewer than 5 clones contained the minor allele. Version 4.03 of the DM reference genome was used for physical positions (Potato Genome Sequencing Consortium 2011; Sharma et al. 2013).

An augmented experimental design was used for field trials at the Hancock Agricultural Research Station (Hancock, Wisconsin), with one plot for each breeding line and three (in 2017) or four (in 2018) plots per check variety. (Several clones besides the checks have more than one data point per environment because they were later determined to be genetic duplicates from the SNP array data and renamed.) Each plot was planted with 15 seed pieces, with 30 cm in-row spacing and 90 cm between rows. The experiments were planted on April 24, 2017, and May 1, 2018. Vine maturity was measured at 100 days after planting (DAP) using a 1 (early) to 9 (late) visual rating of senescence. Plots were harvested 120 days after planting (DAP), and diquat bromide was applied at 14 and 7 days before harvest to promote vine desiccation. Tubers were mechanically harvested into 30 × 45 cm rigid plastic milk crates, run through the standard washing and grading line, and then crated for storage at 12°C with 95% relative humidity.

Skin maturity was measured from images collected within a few days of harvest, following the protocol of Caraza-Harter and Endelman (2020). A representative sample of 7-8 tubers from each plot was photographed on both sides against a black background, using a Photosimile 200 Lightbox and CanonEOS T5i camera. All photos included a MacBeth color card to adjust for lighting and exposure variation. Using the ImageJ software (Schneider et al. 2012), hue and brightness thresholds were used to delineate the skinned surface area for measurement, and each photo was visually inspected to ensure accuracy. Skin maturity is reported as the ratio between skinned area and total surface area, with a log-transformation, *y* = log (*x* + 1), to improve normality of the residuals.

Whole-genome sequencing for two parents from the diallel population (W6511-1R, Villetta Rose) utilized the NovaSeq S1 2x150 flow cell (University of Minnesota Genomics Center).

Reads were aligned with BWA-MEM (Li 2013) to the *CDF1.1_scaffold1389* and *CDF1.3_scaffold1390* alleles from the Atlantic reference genome (Hoopes et al. 2022). The software SAMtools (Li et al. 2009) was used to fix paired reads and remove PCR duplicates. The software BEDtools (Quinlan and Hall 2010) was used to remove reads with fewer than 10 bp on both sides of the transposon insertion site (Gutaker et al. 2019). The software IGV (Robinson et al. 2011) was used to visualize the alignments and count *CDF1* allele read depth. A multiple sequence alignment of *CDF1* alleles from a tetraploid potato pan-genome (Hoopes et al. 2022) was performed using MUSCLE v3.8 (Edgar 2004; Madeira et al. 2019), and the percent identity matrix was used for UPGMA hierarchical clustering with *hclust* in R (R Core Team 2021).

### Statistical analysis

Data were analyzed in R (R Core Team 2021) based on a two-stage approach (Damesa et al. 2017), using ASReml-R v4 (Butler et al. 2018) for REML estimation of variance components. In Stage 1, the best linear unbiased estimate (BLUE) for genotype was computed separately in each year, with a random cofactor for block. The same model, but with genotype as random, was used to estimate the broad-sense heritability on a plot basis as Vg/(Vg+Ve) from the Stage 1 variance components for genetic (Vg) and residual (Ve) variance. Our initial Stage 2 analysis was based on the following model:

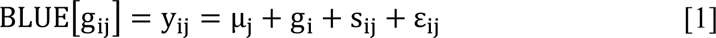

where g_ij_ is the genotypic value for clone *i* in year *j*, μ_j_ is the fixed effect for year *j*, and g_i_ are independent and identically distributed (i.i.d.) normal effects for genotype. The sij random effect follows a multivariate normal distribution with no free variance parameters: the variance-covariance matrix Ω is the direct sum of the variance-covariance matrices of the Stage 1 BLUEs (Damesa et al. 2017). The model residuals ε_ij_ are i.i.d. and represent the genotype x year interaction (Damesa et al. 2017).

The BLUEs from Stage 1 were also used as the response variable for genetic mapping, with year as a fixed effect. Genome-wide association studies utilized the software GWASpoly (Rosyara et al. 2016). Population structure was controlled using a kinship matrix (Yu et al. 2006), calculated with the leave-one-chromosome-out (LOCO) method (Yang et al. 2014). The discovery threshold was based on a significance level of 0.1, corrected for multiple testing by the effective number of markers (Moskvina and Schmidt 2008). Partial R^2^ values were computed based on the change in the likelihood under backward elimination (Sun et al. 2010). Joint linkage mapping used the software PolyOrigin for parental phasing and haplotype construction (Zheng et al. 2021) and diaQTL (Amadeu et al. 2021) for QTL analysis. The discovery threshold was based on a genome-wide significance level of 0.1, established using simulation (Amadeu et al. 2021).

The number of MCMC iterations was determined using the function *set_params* at the recommended settings. For genetic mapping of skin maturity, the Stage 1 BLUEs for vine maturity were used as a covariate.

The genomic partitioning of variance for the Stage 1 BLUEs was based on an extension of the original Stage 2 model (Eq. 1). In the new model, the main effect for genotype (gi) combines fixed effects for significant QTL with random polygenic effects:

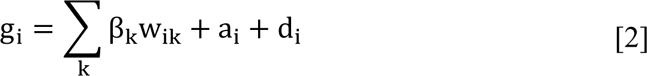

In the above equation, w_ik_ is the (centered) allele dosage for clone *i* at marker *k*, and β_k_ is the fixed effect for marker *k*. The vector of additive values, **a**, follows a multivariate normal distribution with covariance proportional to a genomic relationship matrix (**G**) estimated from markers (VanRaden 2008; Endelman et al. 2018). The vector of digenic dominance values, **d**, is also multivariate normal based on a genomic dominance matrix (Endelman et al. 2018).

Following typical practice for the calculation of heritability, the Stage 1 BLUEs were adjusted for the estimated year effect before computing the proportion of variation for each term based on Legarra (2016). Path coefficients were computed from the proportions of variation to be consistent with the equation of complete determination (Wright 1934; Lynch and Walsh 1998).

## RESULTS

### Vine maturity

Over a two-year period, 586 clones from the US red-skin market type were phenotyped for vine maturity at 100 days after planting. Based on replicated checks, the estimated broad-sense heritability was 0.54 in 2017 and 0.77 in 2018. In a multi-year analysis, the variance of the main effect for genotype was three times the magnitude of the genotype x year interaction (Table 1), so QTL mapping was conducted based on both years.

**Table 1.** Partitioning of variance for vine and skin maturity in a two-year field trial. Vine maturity used a 1 (early) to 9 (late) visual rating, and skin maturity is reported as log(x+1), where x is the skinned area percentage.

|  | Vine Maturity | Skin Maturity |
| --- | --- | --- |
| Year | 0.19 | 0.026 |
| Genotype | 2.61 | 0.156 |
| Genotype x Year | 0.85 | 0.118 |
| Residual | 1.17 | 0.110 |

GWAS identified a QTL on chromosome 5 in the vicinity of *CDF1* that explained 33% of the variation, and a minor QTL on chromosome 6 explained 4% (Table 2, Fig. S1). Figure 2 illustrates the additive relationship between vine maturity and allele dosage at the most significant SNP, PotVar0079081 (49 kb from *CDF1*), with an average maturity decrease of 1.8 per dose of the “B” allele from the SNP array.

**Figure 2.**
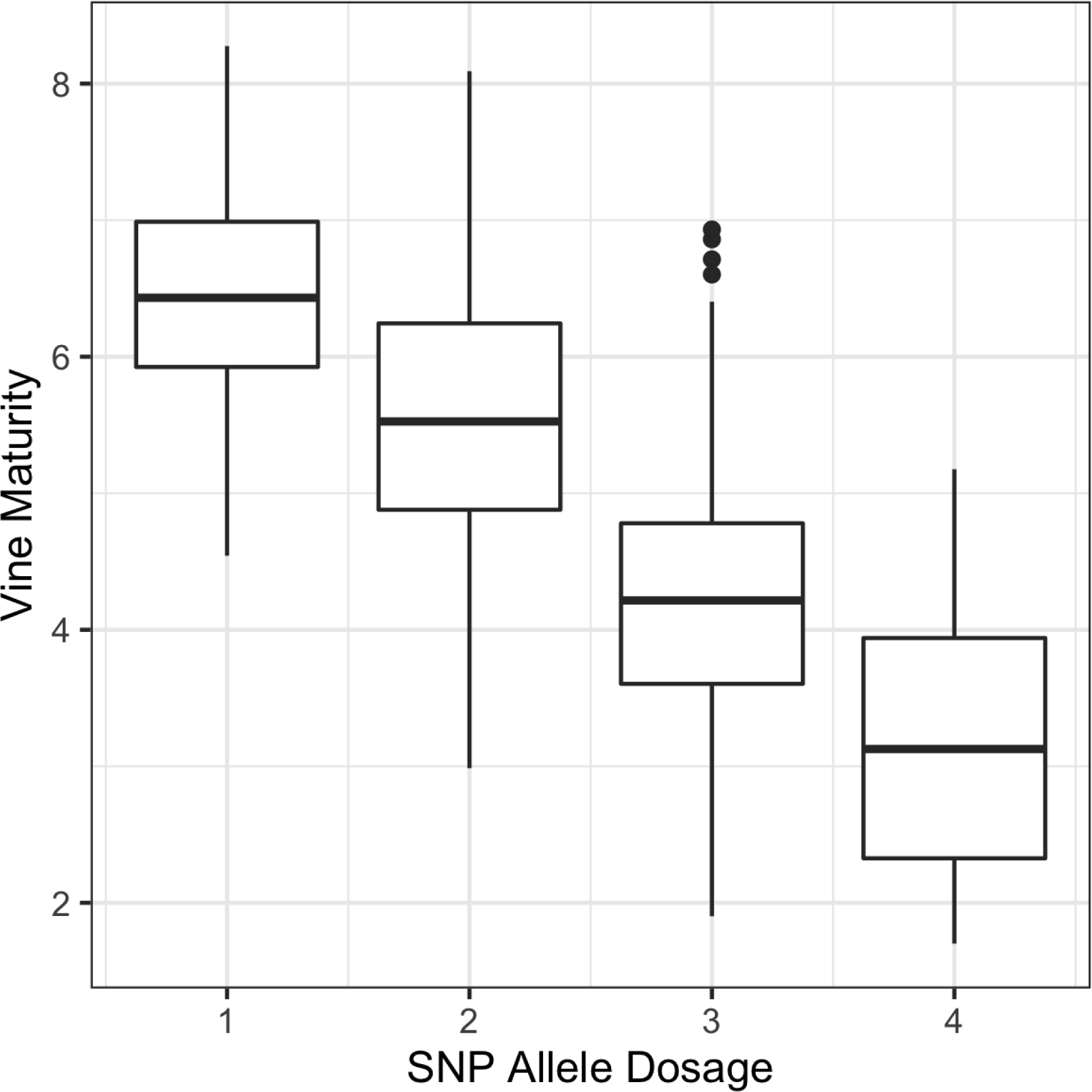
Illustrating the additive relationship between vine maturity and allele dosage at SNP PotVar0079081, which is 49 kb from *CDF1*.

**Table 2.** Genome-wide association study results for maturity.

| Trait | Marker | Chrom | Position (bp) | % Variance Explained |
| --- | --- | --- | --- | --- |
| Vine | PotVar0079081 | 5 | 4,489,481 | 33 |
| Vine | PotVar0090539 | 6 | 48,596,917 | 4 |
| Skin | solcap_snp_c2_14718 | 4 | 86,398,597 | 3 |
| Skin | PotVar0079652 | 5 | 4,548,540 | 3 |

Within the GWAS panel was a half-diallel of 397 clones derived from three parents, named W6511-1R, W9914-1R, and Villetta Rose (Fig. S2). Joint linkage mapping also identified a major QTL near *CDF1* that explained 44% of the variance (Table 3, Fig. S3), and a minor QTL on chromosome 4 explained 5%. Based on the estimated additive effects (Fig. 3), a nearly binary classification of haplotypes into late (positive effect) vs. early (negative effect) was possible. The only exception was haplotype 3 (numbering is arbitrary) in parent W6511-1R, which had an intermediate effect on vine maturity. The phased genotype for PotVar0079081 at the bottom of Fig. 3 shows the “B” allele had perfect linkage with early maturity, excluding the W6511-1R.3 haplotype.

**Figure 3.**
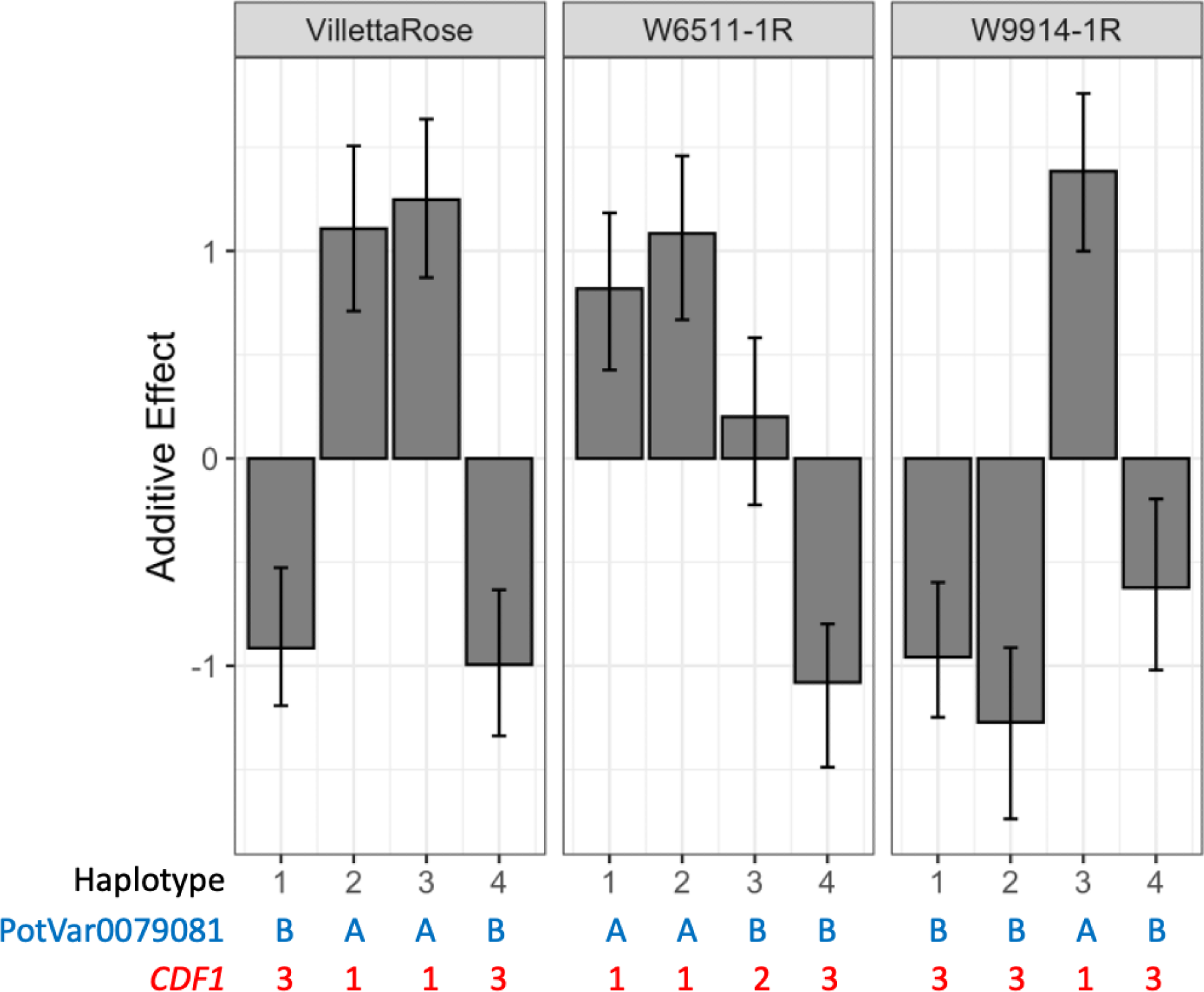
Additive haplotype effects for vine maturity (higher = later) at *CDF1* on chromosome 5. Haplotypes are arbitrarily numbered for the parents of the diallel population (Villetta Rose, W6511-1R, W9914-1R). Error bars represent the 90% CI.

**Table 3.** Joint linkage mapping results for maturity.

| Trait | Marker | Chrom | Position (bp) | % Variance Explained |
| --- | --- | --- | --- | --- |
| Vine | solcap_snp_c2_12921 | 4 | 69,363,089 | 5 |
| Vine | PotVar0079747 | 5 | 4,550,233 | 44 |
| Skin | solcap_snp_c2_4415 | 9 | 8,670,291 | 8 |

Based on previous research summarized in the Introduction, we hypothesized that parental haplotypes with late maturity were wild-type *CDF1.1* alleles, the early haplotypes were *CDF1.3*, and the intermediate haplotype W6511-1R.3 was either *CDF1.2* or *CDF1.4*. Allele counts from the alignment of Illumina short reads against *CDF1* confirmed this hypothesis (Table 4). The tetraploid *CDF1* genotype for Villetta Rose is 1133 and W6511-1R is 1123 (bottom of Fig. 3).

**Table 4.** CDF1 allele read counts from whole-genome sequencing of two parents of the diallel population.

| *CDF1*<br>Allele | Villetta Rose | W6511-1R |
| --- | --- | --- |
| 1 | 36 | 40 |
| 2 | 0 | 26 |
| 3 | 45 | 15 |

Although W9914-1R no longer existed to submit for sequencing, the most likely *CDF1* genotype is 1333.

### Skin maturity

Skin maturity was measured on the same plots as vine maturity, with broad-sense heritability estimates of 0.77 and 0.71 in 2017 and 2018, respectively. The relative magnitude of the genotype x year interaction was larger for skin maturity than vine maturity, but the largest component of variance was still the main effect for genotype (Table 1). The phenotypic correlation between the maturity traits was 0.41 in 2017 and 0.45 in 2018 (Fig. 4).

**Figure 4.**
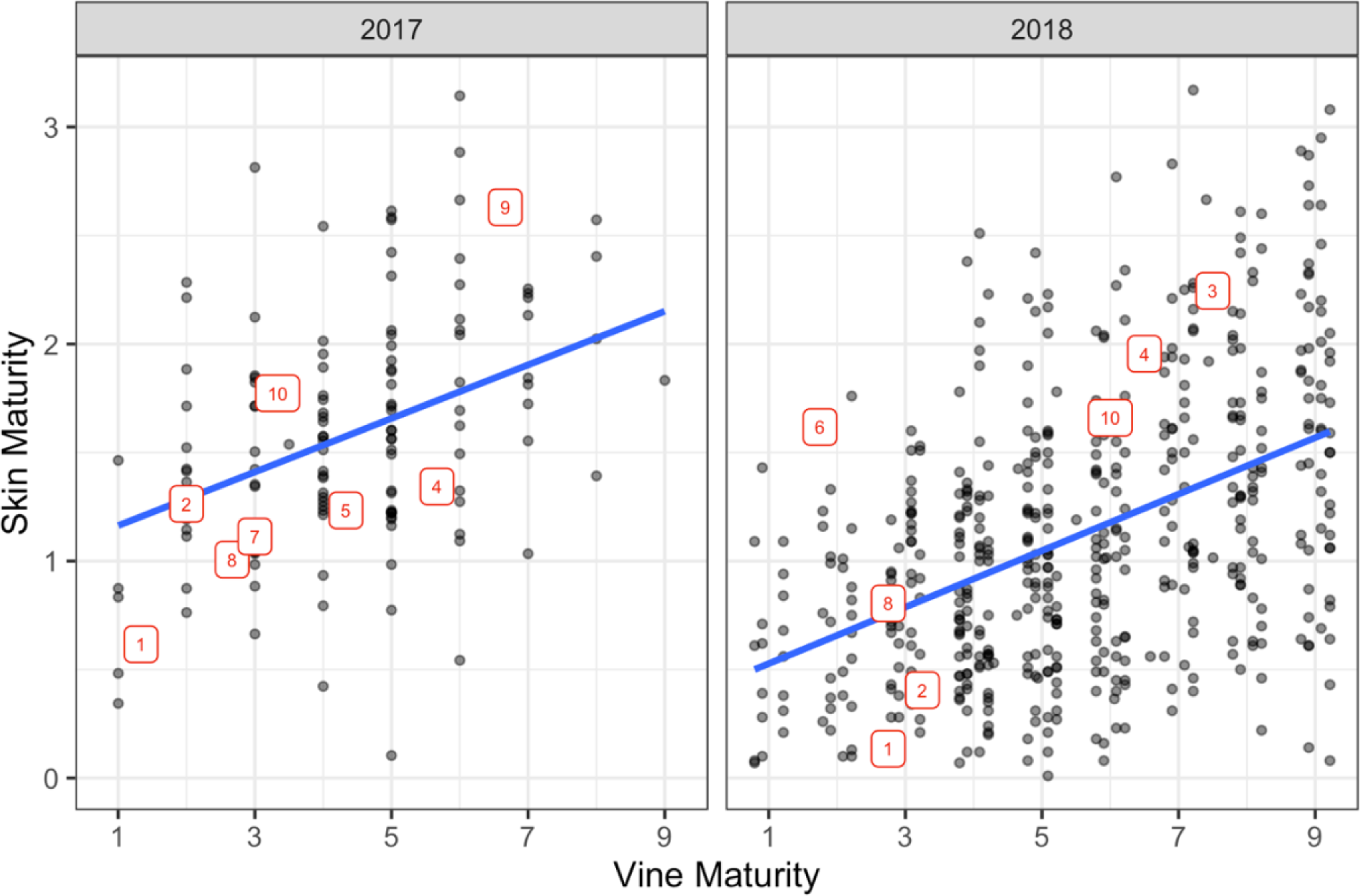
Scatterplot of skin maturity vs. vine maturity from a two-year field trial. Vine maturity used a 1 (early) to 9 (late) visual rating, and skin maturity is reported as log(*x*+1), where *x* is the skinned area percentage. The repeated checks are labeled 1–10 (1 = Red Norland, 2 = Dark Red Norland, 3 = Chieftain, 4 = Red Prairie, 5 = Red Endeavor, 6 = W8893-1R, 7 = W8890-1R, 8 = W10209-2R, 9 = W10209-7R, 10 = Red LaSoda Selection 10). The points are adjusted means for the experimental population, with standard errors (SE) of 0.26 and 0.35 for skin maturity in 2017 and 2018, respectively; for vine maturity, the SE were 1.29 in 2017 and 1.05 in 2018.

One limitation of our experiment was that all plots were harvested at the same time, regardless of their vine maturity. We designate this trait as SMt, to represent skin maturity at a fixed *time*. The actual trait of interest is skin maturity at a fixed physiological *age*, which we designate SMa. Because skin set is promoted by vine senescence, genotypes with earlier VM are expected to have earlier SMt, even if VM and SMa are uncorrelated.

To investigate the genetic architecture of SMa, the response variable SMt was analyzed using VM as a covariate. GWAS identified two minor QTL that each accounted for 3% of the variance (Table 2, Fig. S4), and the QTL on chromosome 5 was inferred to be *CDF1* based on its close proximity (< 10 kb). Joint linkage mapping in the diallel population identified one QTL on chromosome 9, which explained 8% of the variance (Table 3, Fig. S5). From the estimated haplotype effects (Fig. S6), we can infer that one haplotype in W6511-1R carries a unique allele for early skin maturity.

The path diagram in Fig. 5 summarizes the genetic relationships between skin and vine maturity and their explanatory variables. The correlation between VM and SMa due to CDF1 was 0.16 × 0.57 = 0.09. The correlation between VM and SMt had a small component due to the influence of CDF1 on SMa (0.57 × 0.16 × 0.90 = 0.08), but mostly it was due to the direct influence of VM on SMt (0.39), for a total correlation of 0.47. The correlation between the two measures of skin maturity was 0.94, primarily due to the causal influence of SMa on SMt (0.90), but there was also a small contribution from CDF1 through VM (0.57 × 0.16 × 0.39 = 0.04). Genomic heritability can be computed from the diagram as 0.37^2^ + 0.57^2^ = 0.46 for VM, which was larger than the result 0.16^2^ + 0.55^2^ = 0.33 for SMa. The relative importance of additive vs. dominance effects is also depicted in the path diagram. Whereas no dominance variance was estimated for VM, it explained 0.15^2^/(0.15^2^ + 0.55^2^) = 7% of the polygenic effect variance for SMa.

**Figure 5.**
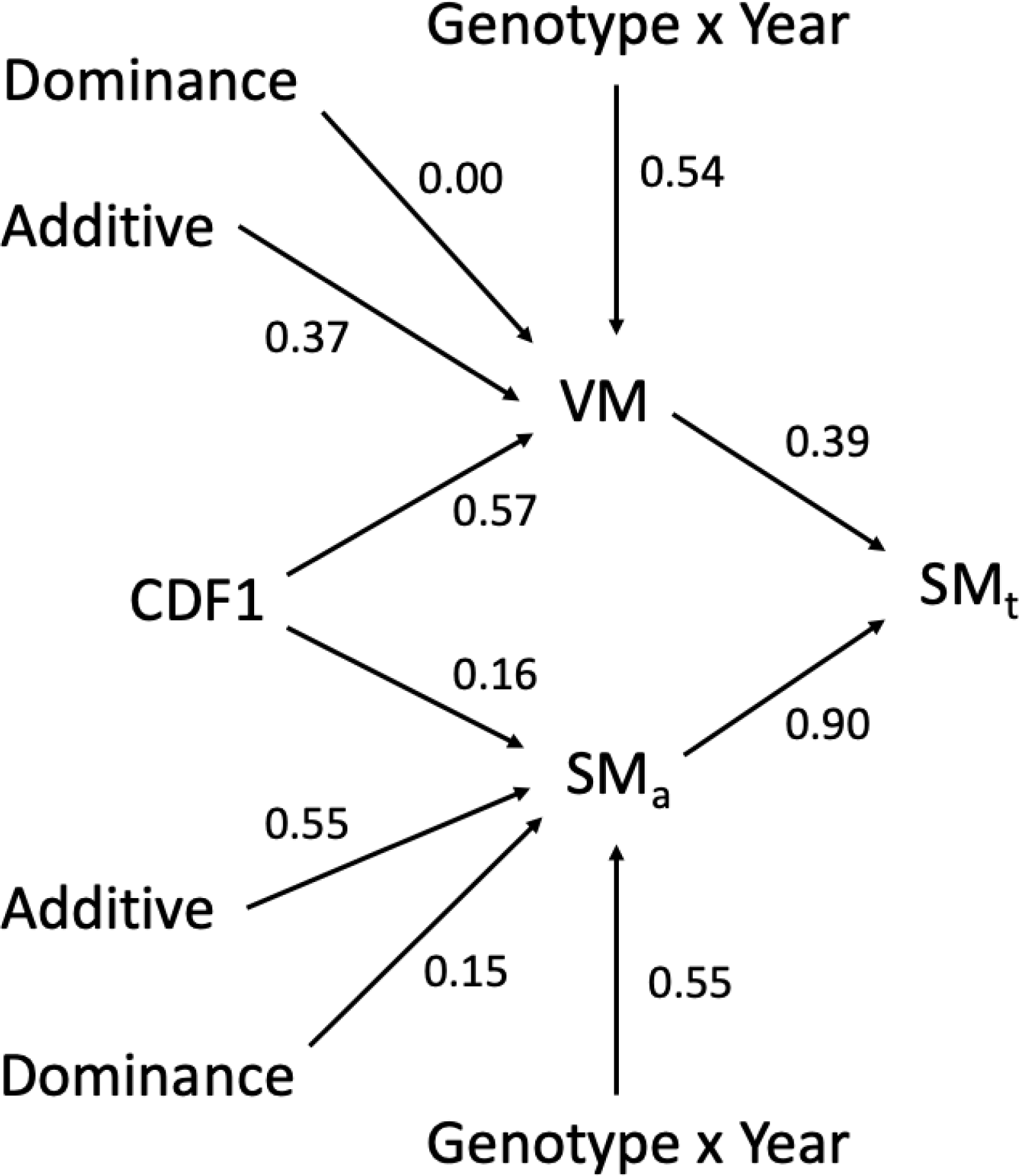
Path diagram of the genetic relationships between potato vine and skin maturity and their explanatory variables. VM = Vine Maturity, SMt = Skin Maturity at a fixed time, SMa = skin maturity at a fixed physiological age.

## DISCUSSION

This is the second publication in which QTL haplotype effects for vine maturity were connected to *CDF1* alleles in a tetraploid mapping population. Hoopes et al. (2022) analyzed a single F1 population with *CDF1* parental genotypes 1111 x 1223, and we have extended the method to a multi-parental population. Our determination of the *CDF1* allelic series 3 > 2 > 1 for early maturity agrees with the results of Hoopes et al. (2022). An additional discovery in our study was the dosage effect of truncated *CDF1* alleles on vine maturity, which could not be observed by Hoopes et al. (2022) based on the parental genotypes. Although not explicitly mentioned by Kloosterman et al. (2013), their supplemental results for vine maturity in a diploid F1 population with *CDF1* parental genotypes 12 x 13 also appear to demonstrate an additive effect for truncated alleles.

The most significant SNP for vine maturity in our GWAS, PotVar0079081, has been reported in other recent publications (Willemsen 2018; Klaassen et al. 2019; Ospina Nieto et al. 2021) that used a GWAS panel with mostly European varieties but also some founders of the US red market type, such as Triumph, Katahdin, and Early Rose. From the pedigree analysis of Vos et al. (2015), we know the SNP predates modern breeding (“pre-1945”). With a physical distance of 49 kb and map distance of 1.5 cM from *CDF1*, PotVar0079081 is not particularly close to the causal locus; there were over 20 SNPs in our dataset closer to *CDF1*. This illustrates a well-known limitation of GWAS panels: the fine-map position of an unobserved causal variant cannot be reliably inferred from marker significance (Schaid et al. 2018).

From the observation that PotVar0079081 had perfect linkage disequilibrium with the *CDF1.2* and *CDF1.3* alleles (as a group) in the parents of our diallel population, a potential explanation for its high significance in association studies can be offered. Genetic distance analysis suggests many different wild-type (*CDF1.1*) alleles existed in the potato gene pool when *CDF1.3* was created by a transposon insertion (Fig. 6). Soon thereafter (on the evolutionary time scale for *CDF1*), the transposon was excised, leaving a 7 bp footprint in *CDF1.2*. If PotVar0079081 arose after diversification of the wild-type alleles, in the lineage where *CDF1.3* arose but before the transposon insertion, then it would be a good proxy for the dosage of late-maturing, wild-type alleles relative to the dosage of truncated alleles in panels with a low frequency of *CDF1.4* (which arose in a separate lineage, see Fig. 6).

**Figure 6.**
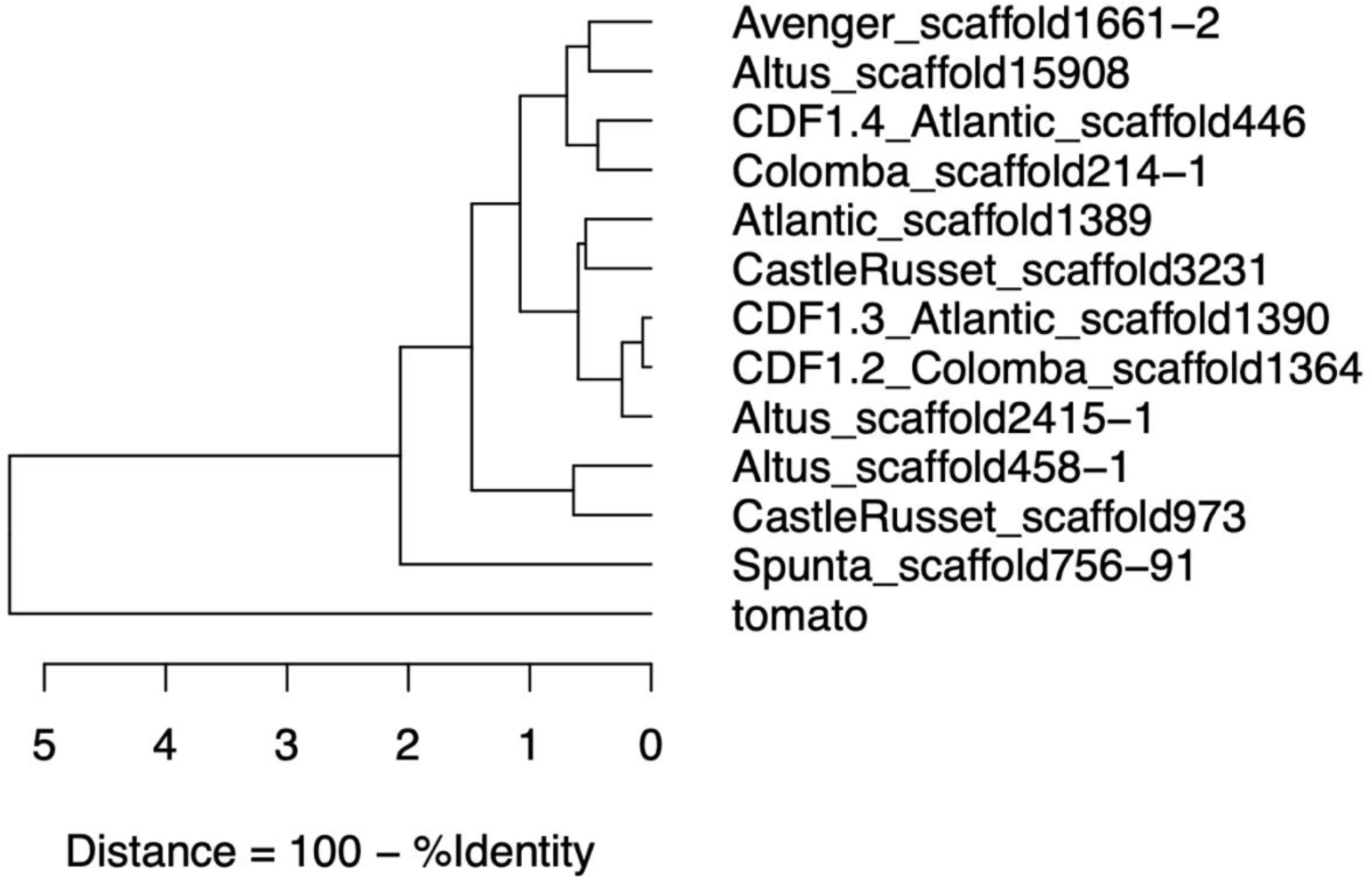
*CDF1* dendrogram based on percent identity from a multiple sequence alignment of 12 potato alleles and one tomato allele. The potato alleles are labeled based on the variety and scaffold identifier from Hoopes et al. (2022). Truncated alleles are labeled *CDF1.2*, *CDF1.3*, *CDF1.4*, and the other nine potato alleles are wild-type *CDF1.1*.

The discovery that CDF1 had a small but statistically significant influence on skin maturity, even after correcting for vine maturity, may be viewed skeptically because there is no known connection between these processes at the molecular level. The result could be an artifact that arises because vine maturity measurements are not a perfect proxy for physiological age.

However, no clear molecular link has been established between CDF1 and vine maturity either, despite the overwhelming evidence for a causal relationship (Collins et al. 1999; Bradshaw et al. 2004; Uitdewilligen et al. 2013; Hoopes et al. 2022). If CDF1 can influence physiological age, this would explain its effects on both vine and skin maturity.

Despite its importance, skin maturity has not been routinely measured in potato breeding as a quantitative trait. One justification may be the belief that selection for early vine maturity (which is routinely measured) will also lead to earlier skin maturity. If the primary goal is improved skin maturity under the same length of the growing season, then the estimated correlation of 0.47 between VM and SMt appears to support this strategy. However, indirect selection is expected to become less efficient over time, as selection against wild-type *CDF1.1* alleles reduces genetic variation for vine maturity. Because the measurement of skin maturity using image analysis is now quite feasible, and because our experiment indicates a genomic heritability of 0.30 even without *CDF1*, the logic for direct selection is compelling.

## Acknowledgments

We thank Peyton Sorensen for assistance with data collection, as well as the staff of the UW Rhinelander and Hancock Agricultural Research Stations.

## Author Contributions

Study design: JBE. Data collection: MCH. Data analysis: MCH, JBE. Manuscript preparation: MCH, JBE.

## Data Availability

The SNP array marker genotypes and phenotype data are available from the Dryad Depository (https://doi.org/10.5061/dryad.3n5tb2rk0). The genotype input file for joint linkage mapping is distributed with the diaQTL package (https://github.com/jendelman/diaQTL). FASTQ files are available from the NCBI Sequence Read Archive under BioProject ID PRJNA823750 (https://www.ncbi.nlm.nih.gov/bioproject/823750).

## Supplemental Figures

**Fig. S1.**
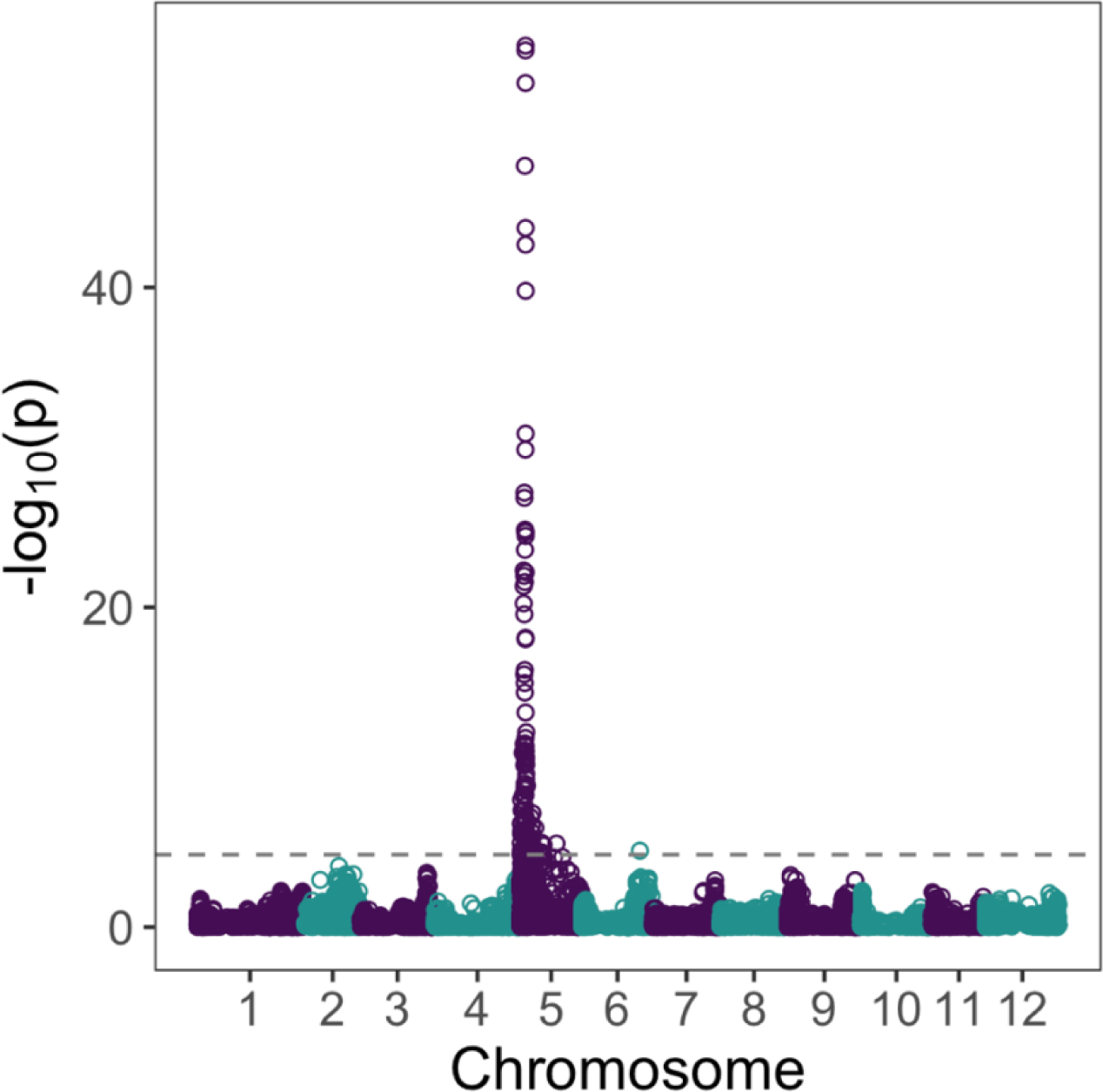
GWAS results for vine maturity with the additive model.

**Fig S2.**
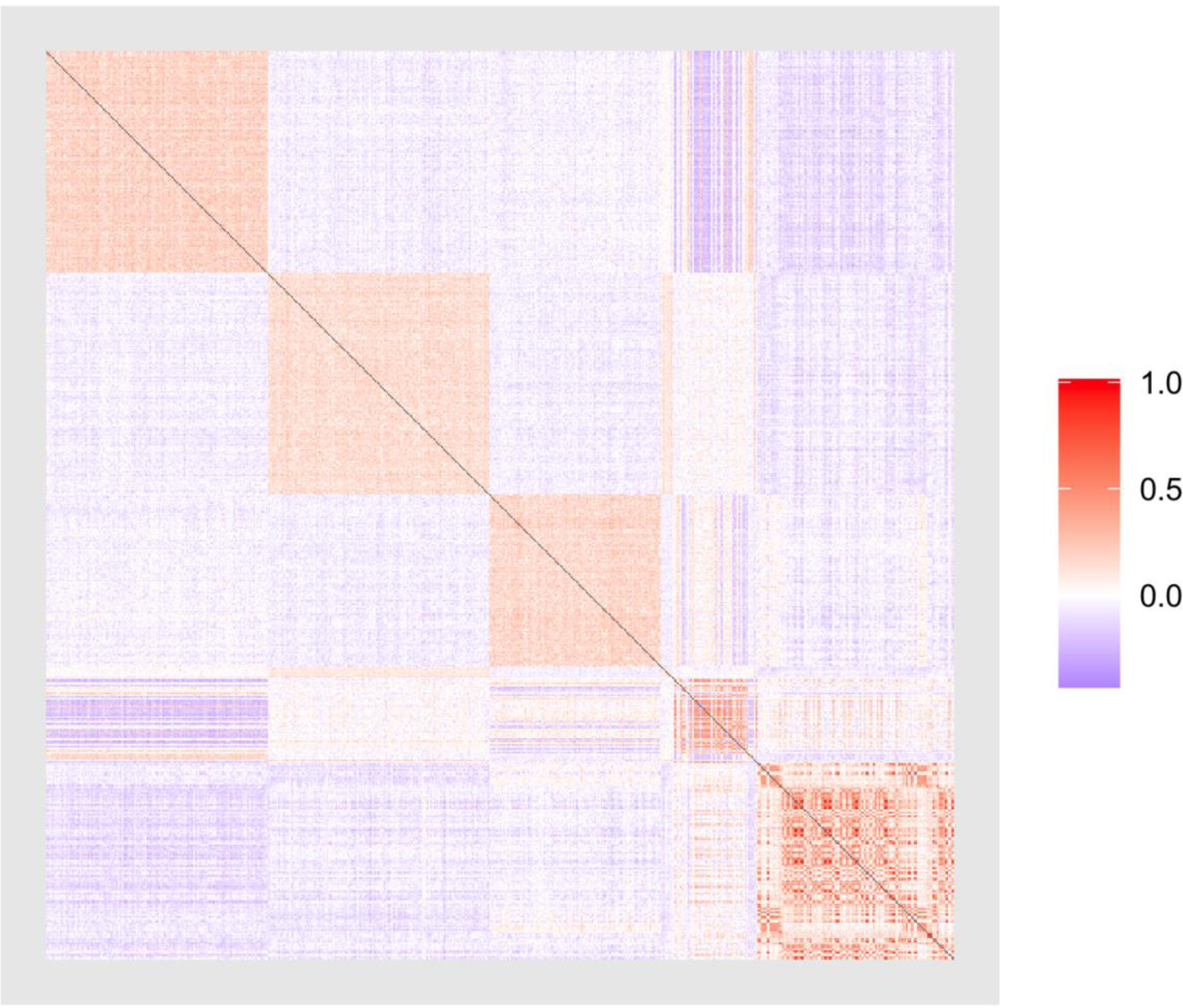
Visualizing population structure in the GWAS panel, based on the additive relationship matrix (G). The first three red squares along the diagonal represent the three F1 populations of the half-diallel. The remainder of the panel consists of F1 populations of varying size from the same breeding program.

**Fig S3.**
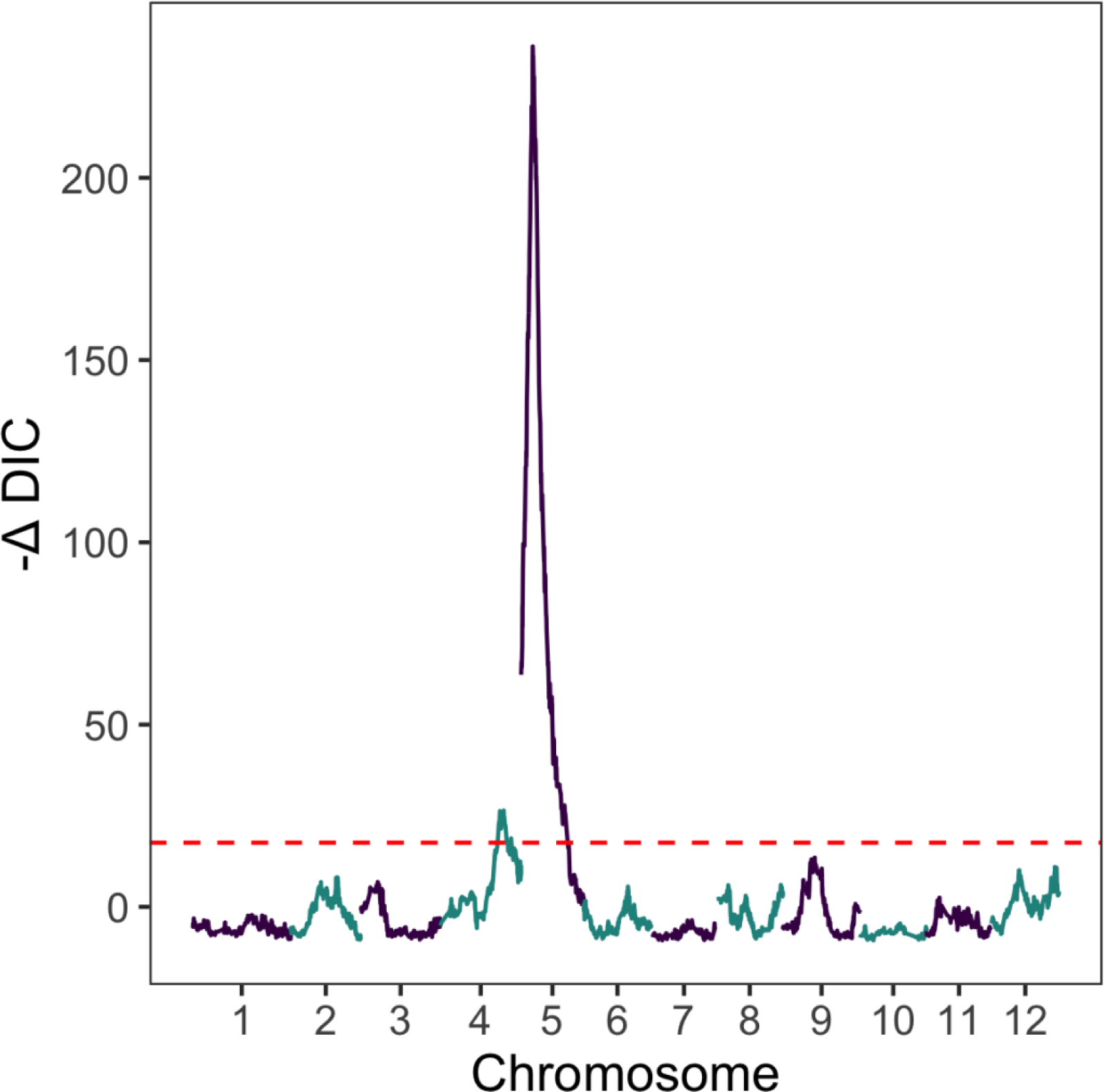
Joint linkage mapping results for vine maturity.

**Fig. S4.**
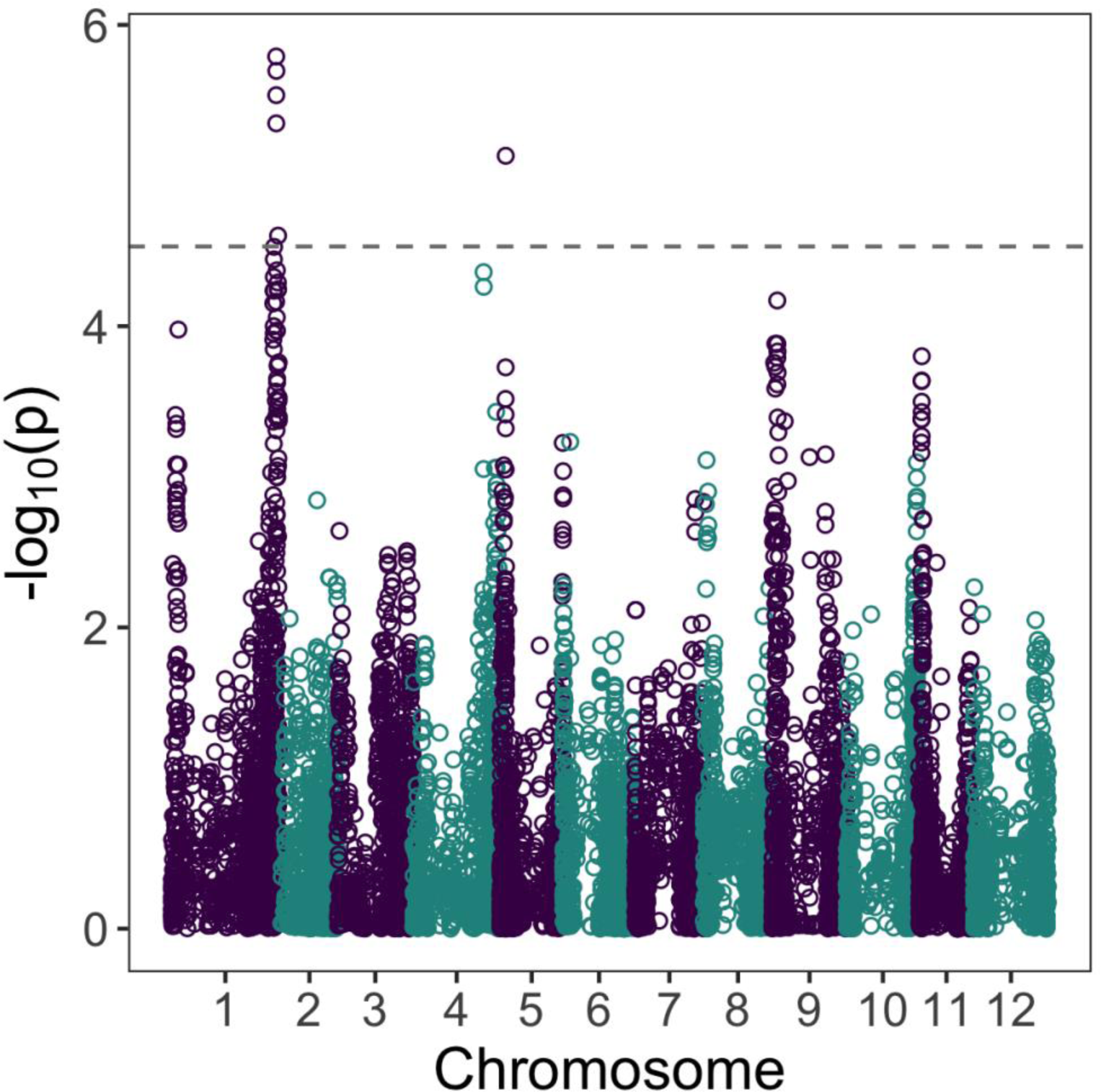
GWAS results for skin maturity with the additive model, using vine maturity as a covariate.

**Fig. S5.**
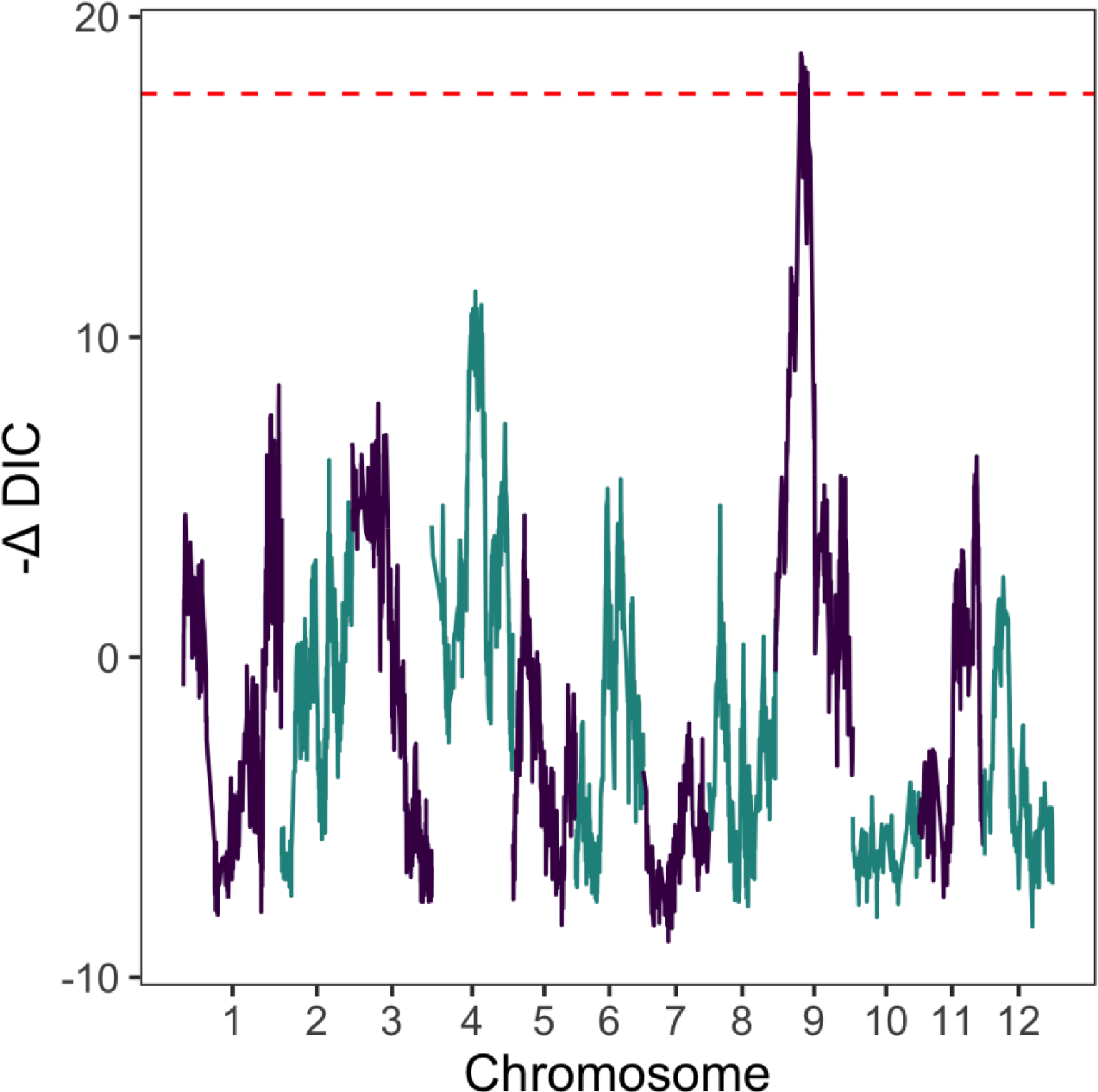
Joint linkage mapping results for skin maturity, using vine maturity as a covariate.

**Fig. S6.**
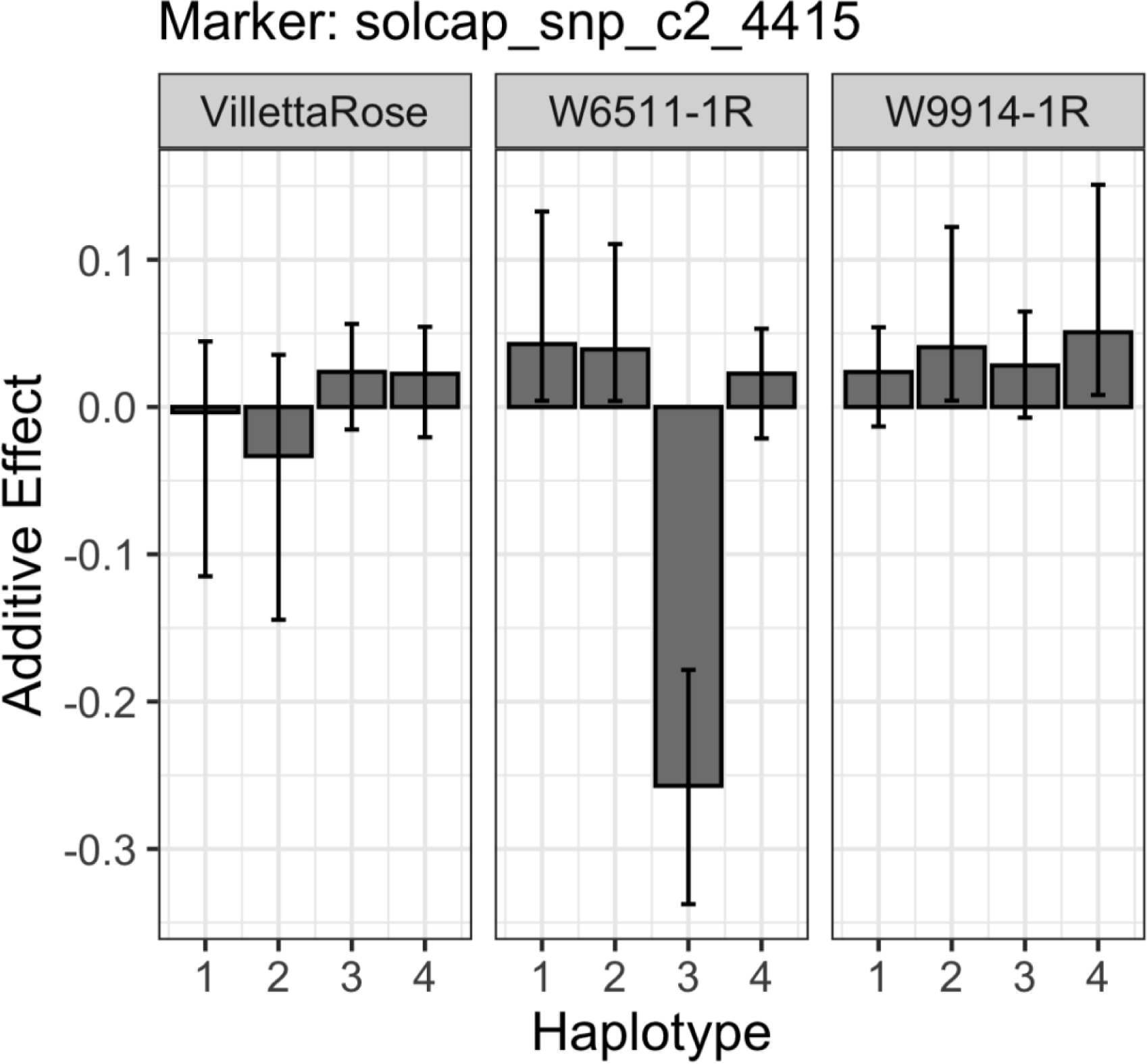
Additive haplotype effects for skin maturity (higher = later) for the QTL on chromosome 9. Haplotypes are arbitrarily numbered for the parents of the diallel population (Villetta Rose, W6511-1R, W9914-1R). Error bars represent the 90% CI.

## REFERENCES

1. Amadeu RR, Muñoz PM, Zheng C, Endelman JB (2021) QTL mapping in outbred tetraploid (and diploid) diallel populations. Genetics 219(3). https://doi.org/10.1093/genetics/iyab124

2. Abelenda JA, Navarro C, Prat S (2011) From the model to the crop: genes controlling tuber formation in potato. Current Opinion in Biotechnology 22: 287–292.

3. Bethke PC, Busse JS (2010) Vine-kill treatment and harvest date have persistent effects on tuber physiology. American Journal of Potato Research 87:299–309. https://doi.org/10.1007/s12230-010-9137-4

4. Bradshaw JE, Pande B, Bryan GJ, Hackett CA, McLean K, Stewart HE, Waugh R (2004) Interval mapping of quantitative trait loci for resistance to late blight [*Phytophthora infestans* (Mont.) de Bary], height and maturity in a tetraploid population of potato (*Solanum tuberosum* subsp. *tuberosum*). Genetics 168:983–995. https://doi.org/10.1534/genetics.104.030056.

5. Bruns HA (2007) A survey of factors involved in crop maturity. Agronomy Journal 101: 60–66.

6. Bussan AJ, Sabba RP, Drilias MJ (2009) Tuber maturation and potato storability: Optimizing skin set, sugars and solids. University of Wisconsin-Extension, Cooperative Extension.

7. Butler D, Cullis B, Gilmour A, Gogel B, Thompson R (2018) ASReml-R Reference Manual Version 4. VSN International Ltd, Hemel Hempstead, HP1 1ES, UK.

8. Caraza-Harter MV, Endelman JB (2020) Image-based phenotyping and genetic analysis of potato skin set and color. Crop Science 60:202–210. https://doi.org/10.1002/csc2.20093

9. Collins A, Milbourne D, Ramsay L, Meyer R, Chatot-Balandras C, Oberhagemann P, De Jong W, Gebhardt C, Bonnel E, Waugh R (1999) QTL for field resistance to late blight in potato are strongly correlated with maturity and vigour. Molecular Breeding 5:387–398.

10. Colwell FJ, Souter J, Bryan GJ, Compton LJ, Boonham N, Prashar A (2021) Development and validation of methodology for estimating potato canopy structure for field crop phenotyping and improved breeding. Frontiers Plant Science 12:612843. https://doi.org/10.3389/fpls.2021.612843.

11. Damesa TM, Möhring J, Worku M, Piepho H-P (2017) One step at a time: Stage-Wise analysis of a series of experiments. Agronomy Journal 109:845–857. https://doi.org/10.2134/agronj2016.07.0395

12. Edgar RC (2004) MUSCLE: a multiple sequence alignment method with reduced time and space complexity. BMC Bioinformatics 5:113. https://doi.org/10.1186/1471-2105-5-113

13. Endelman JB, Schmitz Carley CA, Bethke PC et al. (2018) Genetic variance partitioning and genome-wide prediction with allele dosage information in autotetraploid potato. Genetics 209:77–87. https://doi.org/10.1534/genetics.118.300685

14. Felcher KJ, Coombs JJ, Massa AN, Hansey CN, Hamilton JP, Veilleux RE, Buell CB, Douches DS (2012) Integration of two diploid potato linkage maps with the potato genome sequence. PLoS ONE 7(4):e36347. https://doi.org/10.1371/journal.pone.0036347

15. Gutaker RM, Weiss CL, Ellis D, Anglin NL, Knapp S, Fernández-Alonso JL, Prat S, Burbano HA (2019) The origins and adaptation of European potatoes reconstructed from historical genomes. Nature Ecol Evol 3:1093–1101. https://doi.org/10.1038/s41559-019-0921-3

16. Halderson JL, Henning RC (1993) Measurements for determining potato tuber maturity. American Potato Journal 70:131–141.

17. Hoopes G, Meng X, Hamilton JP et al. (2022) Phased, chromosome-scale genome assemblies of tetraploid potato reveal a complex genome, transcriptome, and proteome landscape underpinning phenotypic diversity. Molecular Plant 15:520–536. https://doi.org/10.1016/j.molp.2022.01.003

18. Klaassen MT, Willemsen JH, Vos PG, Visser RG, Van Eck HJ, Maliepaard C, Trindade LM (2019) Genome-wide association analysis in tetraploid potato reveals four QTLs for protein content. Molecular Breeding 39. https://doi.org/10.1007/s11032-019-1070-8

19. Kloosterman B, Abelenda JA, Carretero Gomez MM, Oortwijn M, de Boer JM, Kowitwanich K, Horvath BM, van Eck HJ, Smaczniak C, Prat S, Visser RGF, Bachem CWB (2013) Naturally occurring allele diversity allows potato cultivation in northern latitudes. Nature 495:246–250. https://doi.org/10.1038/nature11912

20. Legarra A (2016) Comparing estimates of genetic variance across different relationship models. Theor. Pop. Biol. 107:26–30. https://doi.org/10.1016/j.tpb.2015.08.005

21. Li H (2013) Aligning sequence reads, clone sequences and assembly contigs with BWA-MEM. arXiv:1303.3997v2 [q-bio.GN]

22. Li H, Handsaker B, Wysoker A et al. (2009) The sequence alignment/map format and SAMtools. Bioinformatics 25:2078–2079. https://doi.org/10.1093/bioinformatics/btp352

23. Lulai EC, Orr PH (1993) Determining the feasibility of measuring genotypic differences in skin- set. American Journal of Potato Research 70: 599–609.

24. Lulai EC, Freeman TP (2001) The importance of phellogen cells and their structural characteristics in susceptibility and resistance to excoriation in immature and mature potato tuber (*Solanum tuberosum* L.) periderm. Annals of Botany 88:555–561. https://doi.org/10.1006/anbo.2001.1497

25. Lynch M, Walsh B (1998) *Genetics and Analysis of Quantitative Traits*. Sinauer Associates, Sunderland, MA.

26. Madeira F, Park YM, Lee J, Buso N, Gur T, Madhusoodanan N, Basutkar P, Tivey ARN, Potter SC, Finn RD, Lopez R (2019) The EMBL-EBI search and sequence analysis tools APIs in 2019. Nucleic Acids Research 47(W1):W636–W641. https://doi.org/10.1093/nar/gkz268

27. Moskvina V, Schmidt KM (2008) On multiple-testing correction in genome-wide association studies. Genetic Epidemiology 32:567–573. https://doi.org/10.1002/gepi.20331

28. Murphy HJ (1968) Potato vine killing. American Potato Journal 45: 472–478.

29. Navarro C, Abelenda JA, Cruz-Oró E, Cuéllar CA, Tamaki S, Silva J, Shimamoto K, Prat S (2011) Control of flowering and storage organ formation in potato by FLOWERING LOCUS T. Nature 478:119–122. https://doi.org/10.138/nature10431

30. Neubauer JD, Lulai EC, Thompson AL, Suttle JC, Bolton MD, Campbell LG (2013) Molecular and cytological aspects of native periderm maturation in potato tubers. Journal of Plant Physiology 170:413–423. https://doi.org/10.1016/j.jplph.2012.10.008

31. Ospina Nieto CA, Lammerts van Bueren ET, Allefs S, Vos PG, van der Linden G, Maliepaard CA, Struik PC (2021) Association mapping of physiological and morphological traits related to crop development under contrasting nitrogen inputs in a diverse set of potato cultivars. Plants 10:1727. https://doi.org/10.3390/plants10081727

32. Potato Genome Sequencing Consortium (2011) Genome sequence and analysis of the tuber crop potato. Nature 475:189–195. https://doi.org/10.1038/nature10158

33. Quinlan AR, Hall IM (2010) BEDTools: a flexible suite of utilities for comparing genomic features. Bioinformatics 26:841–842. https://doi.org/10.1093/bioinformatics/btq033

34. R Core Team (2021) R: A language and environment for statistical computing. R Foundation for Statistical Computing, Vienna, Austria.

35. Ramírez Gonzales L, Shi L, Bergonzi SB, Oortwijn M, Franco-Zorrilla JM, Solano-Tavira R, Visser RGF, Abelenda JA, Bachem CWB (2021) Potato CYCLING DOF FACTOR 1 and its lncRNA counterpart *StFLORE* link tuber development and drought response. The Plant Journal 105:855–869. https://doi.org/10.1111/tpj.15093

36. Reeve RM, Hautala E, Weaver ML (1969) Anatomy and compositional variation within potatoes I. Developmental histology of the tuber. American Journal of Potato Research 46:361–373.

37. Robinson JT, Thorvaldsdóttir H, Winckler W et al. (2011) Integrative Genomics Viewer. Nature Biotech. 29:24–26.

38. Rosyara UR, De Jong WS, Douches DS, Endelman JB (2016) Software for genome-wide association studies in autopolyploids and its application to potato. Plant Genome 9:2. https://doi.org/10.3835/plantgenome2015.08.0073

39. Sankaran S, Khot LR, Espinoza CZ, Jarolmasjed S, Sathuvalli VR, Vandemark GJ, Miklas PN, Carter AH, Pumphrey MO, Knowles NR, Pavek MJ (2015) Low-altitude, high resolution aerial imaging systems for row and field crop phenotyping: A review. European Journal Agronomy 70:112–123. https://doi.org/10.1016/j.eja.2015.07.004.

40. Sawa M, Nusinow DA, Kay SA, Imaizumi T (2007) FKF1 and GIGANTEA complex formation is required for day-length measurement in *Arabidopsis*. Science 318:261–265.

41. Schaid DJ, Chen W, Larson NB (2018) From genome-wide associations to candidate causal variants by statistical fine-mapping. Nature Reviews Genetics 19:491–504. https://doi.org/10.1038/s41576-018-0016-z.

42. Schneider C, Rasband W, Eliceiri K (2012) NIH Image to ImageJ: 25 years of image analysis. Nature Methods 9.

43. Searle SR, Casella G, McCulloch CE (1992) Variance components. John Wiley & Sons, Hoboken, NJ.

44. Sharma SK, Bolser D, de Boer J et al (2013) Construction of reference chromosome-scale pseudomolecules for potato: integrating the potato genome with genetic and physical maps. G3 3:2031–2047. https://doi.org/10.1534/g3.113.007153

45. Sowokinos JR (1978) Relationship of harvest sucrose content to processing maturity and storage life of potatoes. American Potato Journal 55:333–344.

46. Struik PC and Wiersema SG (1999) Seed potato technology. Wageningen Academic Publishers, Wageningen.

47. Sun G, Zhu C, Kramer MH, Yang S-S, Piepho H-P, Yu J (2010) Variation explained in mixed-model association mapping. Heredity 105:333–340.

48. Uitdewilligen JGAML, Wolters AMA, D’hoop BB, Borm TJA, Visser RGF, van Eck HJ (2013) A next-generation sequencing method for genotyping-by-sequencing of highly heterozygous autotetraploid potato. PLoS ONE 8:e62355.

49. VanRaden PM (2008) Efficient methods to compute genomic predictions. J. Dairy Sci. 91:4414– 4423.

50. Voorrips RE, Gort G, Vosman B (2011) Genotype calling in tetraploid species from bi-allelic marker data using mixture models. BMC Bioinformatics 12:172. https://doi.org/10.1186/1471-2105-12-172

51. Vos PG, Uitdewilligen JGAML, Voorrips RE, Visser RGF, van Eck HJ (2015) Development and analysis of a 20K SNP array for potato (*Solanum tuberosum*): an insight into the breeding history. Theor Appl Genet 128:2387–2401. https://doi.org/10.1007/s00122-015-2593-y

52. Vulavala VKR, Fogelman E, Faigenboim A, Shoseyov O, Ginzberg I (2019) The transcriptome of potato tuber phellogen reveals cellular functions of cork cambium and genes involved in periderm formation and maturation. Scientific Reports 9:10216. https://doi.org/10.1038/s41598-019-46681-z

53. Willemsen JH (2018) The identification of allelic variation in potato. Wageningen University, Wageningen, the Netherlands.

54. Wright S (1934) The method of path coefficients. Ann. Math. Statist. 5:161–215. https://doi.org/10.1214/aoms/1177732676

55. Yang J, Zaitlen NA, Goddard ME, Visscher PM, Price AL (2014) Advantages and pitfalls in the application of mixed-model association methods. Nature Genetics 467:100–106. https://doi.org/10.1038/ng.2876

56. Yu J, Pressoir G, Briggs WH, Vroh Bi I, Yamasaki M, Doebley JF, McMullen MD, Gaut BS, Nielsen D, Holland JB, Kresovich S, Buckler ES (2006) A unified mixed-model method for association mapping that accounts for multiple levels of relatedness. Nature Genetics 38:203–208. https://doi.org/10.1038/ng1702

57. Zheng C, Amadeu RR, Muñoz PM, Endelman JB (2021) Haplotype reconstruction in connected tetraploid F1 populations. Genetics 219(2). https://doi.org/10.1093/genetics/iyab106

58. Zych K, Gort G, Maliepaard CA, Jansen RC, Voorrips RE (2019) FitTetra 2.0: improved genotype calling for tetraploids with multiple population and parental data support. BMC Bioinformatics 20:148. https://doi.org/10.1186/s12859-019-2703-y

